# Myosin-Binding Protein C Stabilizes, But Is Not the Sole Determinant of SRX Myosin in Cardiac Muscle

**DOI:** 10.1101/2022.10.03.510676

**Authors:** Shane Nelson, Samantha Beck Previs, Sakthivel Sadayappan, Carl Tong, David M. Warshaw

## Abstract

The myosin Super Relaxed (SRX) state is central to striated muscle metabolic and functional regulation. In skeletal muscle, SRX myosin are predominantly colocalized with Myosin-Binding Protein C (MyBP-C) in the C-zone, proximal to the sarcomere center. To define how MyBP-C and its specific domains contribute to stabilizing the SRX state in cardiac muscle, we took advantage of transgenic MyBP-C null mice and those expressing MyBP-C with a 271 residue N-terminal truncation. Utilizing super-resolution microscopy, we determined the lifetime and sub-sarcomeric location of individual fluorescent-ATP turnover events within isolated cardiac myofibrils. The proportion of SRX myosin was highest in the C-zone (71±6%) and lower in the D-zone (45±10%) which lies farther from the sarcomere center and lacks MyBP-C, suggesting a possible role for MyBP-C in stabilizing the SRX state within the C-zone. However, myofibrils from MyBP-C null mice demonstrated an ~40% SRX reduction, not only within the now MyBP-C-free C-zone (49±9% SRX), but also within the D-zone (22±5% SRX). These data suggest that the influence of MyBP-C on the SRX state is not limited to the C-zone, but extends along the thick filament. Interestingly, myofibrils with N-terminal truncated MyBP-C had an SRX content and spatial gradient similar to the MyBP-C null, indicating that the N terminus of cardiac MyBP-C is necessary for MyBP-C’s role in establishing SRX along the entire thick filament. Given that SRX myosin are enriched in the C-zone, even in the absence of MyBP-C or its N-terminus, an inherent bias must exist in the structure of the thick filament to stabilize the SRX state. One candidate may be a differential in the super repeats of titin that interact with MyBP-C and myosin and which template the thick filament.

## Introduction

Striated muscle (i.e., skeletal and cardiac) activation requires calcium binding to the thin filament regulatory complex (i.e., troponin and tropomyosin) which shifts the position of the complex to allow force-generating myosin motors access to actin binding sites along the thin filament (Fig. 1A) (McKillop and Geeves, 1993; Lehman et al., 1995). Recently, a myosin thick filament-based regulatory mechanism has been proposed, wherein myosin heads can adopt an energetically economical “super relaxed” (SRX) state that is also effectively incapable of binding to the thin filament (Stewart et al., 2010; Fig. 1A). The SRX state is biochemically distinct with a very low ATP hydrolysis rate of ~0.005s^−1^, or about 1 ATP molecule every 200s. This rate is 5-fold lower than that of “disordered relaxed” (DRX) myosin that can bind to the thin filament and generate force once muscle is calcium activated (Stewart et al., 2010). The SRX biochemical state may be inherent to each of myosin’s two heads (Anderson et al., 2018; Rhode et al., 2018; Chu et al., 2021; Walklate et al., 2022), but is further stabilized when the two heads are folded back and interact with each other and the myosin tail (i.e., Interacting Heads Motif or IHM) (Yang et al., 2020). This structural motif has been observed in high resolution, 3D electron microscopic reconstructions of native cardiac thick filaments (Al Khayat et al., 2013; Zoghbi et al., 2008). Interestingly, a population of cardiac myosin remains sequestered in the mechanically inactive SRX state during calcium activation of the sarcomere (Hooijman et al., 2011), possibly serving as a “reserve pool” that can be recruited from, in response to physiological demands. Importantly, cardiac myosin mutations associated with hypertrophic cardiomyopathy (HCM) have been linked to a reduced proportion of myosin in the SRX state (McNamara et al., 2017; Toepfer et al., 2019). If so, this may explain the hypercontractility of hearts in hypertrophic cardiomyopathy (HCM) patients and support the therapeutic basis of Mavacamten, an FDA approved small molecule that stabilizes the SRX state as a means of restoring the normal proportion of SRX myosin and thus normal contractility (Green et al., 2016; Anderson et al., 2018).

**Figure 1.**
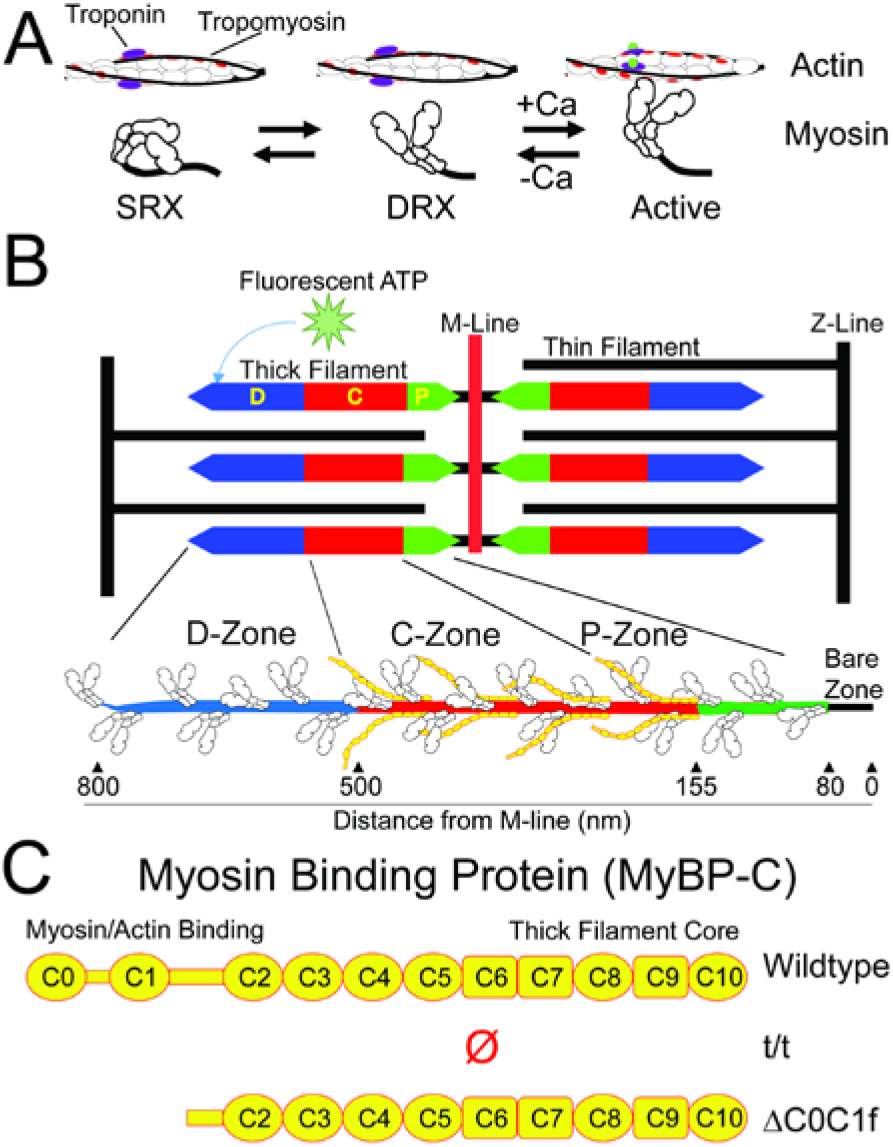
Structure and Regulation of Myosin, cMyBP-C and the Sarcomere. A) Myosin’s biochemical states. B) Cartoon of sarcomere, showing thick and thin filaments, delineation of C-, D-, and P-, and bare zones, as well as myosin and cMyBP-C. C) Domain structure of cMyBP-C, truncated variant (ΔC0C1f) and null (t/t).

As the SRX state may represent a key mechanism for both cardiac energetic and contractile regulation, defining the physiological and structural determinants of the SRX state is critically important. We recently described a super-resolution imaging approach in rat soleus myofibrils to visualize single, fluorescently-labeled ATP being hydrolyzed by myosin within individual sarcomeres (Nelson et al., 2020; Fig. 1B). Interestingly, SRX myosin were not uniformly distributed along the thick filament, but were predominantly found in the C-zone where myosin-binding protein C (MyBP-C) only exists. The flanking P- and D-zones are devoid of MyBP-C, and demonstrated much lower frequencies of SRX myosin (Nelson et al., 2020). These data and that of others implicate MyBP-C in stabilizing the SRX state (McNamara et al., 2017; Toepfer et al., 2019).

MyBP-C is a ~140KDa protein, encoded by 3 orthologous and muscle fiber-type specific genes (fast-type, slow-type, and cardiac). The cardiac isoform, cMyBP-C, is composed of 11 Ig-like and Fn3-like domains denoted as C0 through C10 (Fig. 1C). cMyBP-C is anchored to the thick filament backbone through its C-terminal domains (C8-C10; Lee et al., 2015; Gilbert et al., 1996), whereas the N-terminal domains (C0-C2) are critical to cMyBP-C’s capacity to modulate cardiac contractility through interactions with the myosin head region and/or actin (van Dijk et al., 2014; Fig. 1C). Mutations in cMyBP-C are the leading cause of HCM (Walsh et al., 2017) and are also associated with reductions in the SRX state (McNamara et al., 2017), further emphasizing a potential role for cMyBP-C in regulation of the SRX state.

To define cMyBP-C’s role in stabilizing the SRX state and the involvement of its N terminus, we took advantage of genetically modified mouse cardiac muscle that was either devoid of cMyBP-C (McConnell et al., 1999) or that expressed cMyBP-C lacking the first 29kD of the N terminus (Lynch et al., 2021). Here, we applied the super-resolution imaging of single fluorescently-labeled ATP hydrolysis events in myofibrils from these mouse models to define the spatial distribution of SRX myosin in the presence, absence, and N-terminal truncation of cMyBP-C.

## Materials and Methods

### Mouse models

All experiments using animals detailed in this work were approved by the Institutional Animal Care and Use Committees at the University of Vermont and University of Cincinnati and followed the policies of the Guide for the Use and Care of Laboratory Animals published by the National Institutes of Health. cMyBP-C null (t/t) mice were generated by insertion of a 2kb neomycin resistance gene into exon 30 of MYBPC3 (McConnell et al., 1999). While truncated cMyBP-C (MyBP-C^ΔC0C1f^) were generated by cardiomyocyte-specific replacement of endogenous cMyBP-C with an equivalent cDNA lacking the first 813 nucleotides, as previously described in Lynch et al. (2021). For wildtype controls (WT), frozen hearts from FVB mice were ordered from Jackson Labs (Bar Harbor, ME)

### Preparation of mouse cardiac myofibrils

Mouse cardiac ventricular myofibrils were prepared from flash-frozen mouse hearts following a protocol modified from that described in Creed and Tong (2021). Briefly, the apex of the frozen heart was removed using a razorblade and rinsed twice to remove blood in 1mL of K60 buffer (60mM KCl, 2mM MgCl2, 20mM MOPS, pH 7.4) with 30mM BDM, 1mM EGTA and protease inhibitor cocktail (P8340, Sigma Aldrich, St Louis MO). Tissue was then placed in 1mL K60 buffer and homogenized using a “Tissue Tearor” homogenizer (Biospec Products, Bartlesville, OK) operated for two rounds at 21,000 RPM for 30s each. Tissue was collected by centrifugation at 1000xg for 10min at 4°C and then permeablized in K60 buffer with 1% v/v Triton X-100, 1mM EGTA and protease inhibitor cocktail for 30min. Myofibrils were collected by centrifugation at 1000xg for 5min at 4°C and resuspended in 1mL of K60 buffer, followed by 3 more washes with K60 buffer with 1mg/mL BSA.

### Imaging and antibody labeling conditions

Fluorescence imaging was performed as described (Nelson et al., 2020). To identify the sarcomere center (M-line) where myomesin resides, prior to imaging, each myofibril preparation was incubated with anti-myomesin antibody (Cat #EPR17322-9, Abcam, Cambridge UK) at a 1:5000 final dilution for 15 minutes on ice, followed by Alexa 647 goat-anti-mouse IgG (Cat #A21237, Life Technologies, Carlsbad CA) at a final concentration of 1:5000. After 5 minutes on ice, labeled myofibrils were flowed into flowcells which were made using plasma-cleaned glass. The flowcell was then incubated at room temperature for 20 minutes to allow myofibrils to adhere to the coverglass surface. Finally, 100μL of Relaxing Buffer (120mM KOAc, 5mM K-phosphate, 4mM EGTA, 4mM MgCl2, 50mM MOPS, 4mM ATP, 10nM BODIPY-ATP (Cat #A12410, ThermoFisher, Waltham MA), pH 6.8) was flowed into the flowcell and samples were imaged using a custom-built dual camera super-resolution microscope, described previously (Nelson et al., 2020) for 50 minutes at 10FPS. From each myofibril preparation 3-4 recordings were made, with each flowcell being used for only a single imaging session.

### Fluorescent ATP binding event analysis

The lifetime and position of each BODIPY-ATP binding event was measured separately to ensure accuracy. BODIPY-ATP binding events (green channel) were manually documented in image stacks using ImageJ (Schindelin et al., 2012). First, image stacks were integrated to 1FPS using the “grouped Z-project” function. Image stacks were then converted into a stack of kymographs using the “Reslice” function and events were documented using rectangular selections and the “overlay” function. Event lifetimes were determined manually from kymographs (Fig. 2A) to account for visually apparent blinking behavior, counting only events with visually apparent beginning and endings (i.e., not cut off by the beginning or end of the recording). Super-resolution position information was determined using the ImageJ plugin “D.O.M.” (Katrukha, 2020). Finally, BODIPY-ATP binding lifetimes (from kymograph analysis) were connected to sub-pixel localizations in the sarcomere, as the kymograph X, Y, and Slice coordinates can be directly related to Timestamp, X, and Y coordinates (respectively) in the DOM output files.

**Figure 2.**
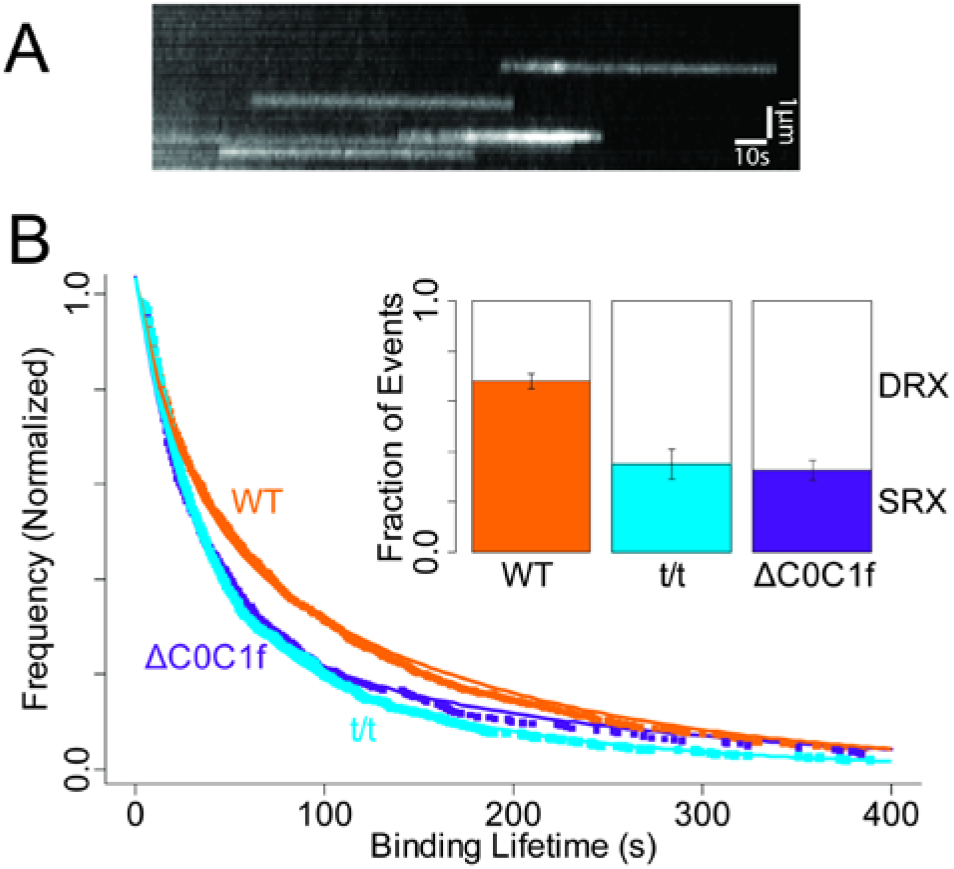
ATP Binding Events and Overall Lifetime Distributions. A) Kymograph showing multiple fluorescent nucleotide binding events (white streaks). B) Overall lifetime distributions for WT, t/t, and ΔC0C1f transgenic. Inset shows proportions of SRX and DRX events.

As previously described (Nelson et al., 2020), overall fluorescence event lifetime distributions for each mouse strain (Fig. 2B) were fit with a double exponential model that accounts for a lack of the shortest event lifetimes in our experimental distributions:

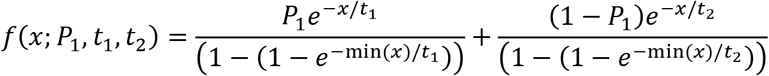

*P_1_* is the proportion of “relaxed” events, *t_1_* is the time constant for the relaxed DRX population, and *t_2_* is the time constant of the SRX population. Lifetime distributions were fit by direct optimization of the log-likelihood estimates as implemented in the “fitdistr” function included in the “MASS” library for the statistical programming language “R” (Venables, 2002) using the “L-BFGS-B” method (Byrd et al., 1995). For convergence reasons when fitting zone-specific lifetime distributions, *t_1_* and *t_2_* were constrained to the values obtained from the overall lifetime fits.

Binding event lifetime distributions were compared using a Kolmogorov-Smirnov test, while sarcomere lengths were compared using a Student’s t-test.

To localize M-line positions, myomesin-labeled (red channel) images were integrated in blocks of 1000 frames using the “Grouped Z-project” function in ImageJ. Each M-line was tracked using the ImageJ plugin “FilamentJ” (Smith et al., 2010), using default settings, except for “Deform Iterations”, which was set to 500. Finally, for each BODIPY-ATP binding event, sub-sarcomeric location was determined as the distance to the nearest M-line at the corresponding time point, after correction for channel registration (described below).

Spatial offsets between red and green imaging channels were calculated by imaging multicolor-fluorescent beads (Tetraspeck, ThermoFisher) and analysis using the Particle Image Velocimetry plugin for ImageJ (Tseng et al., 2012), which calculates relative offsets between the two images by maximizing cross-correlation between corresponding regions in the two images.

## Results

### Single Fluorescent ATP Turnover Events

When myofibrils from WT mouse hearts were imaged under relaxing conditions (no calcium, 4mM ATP) with the addition of 10nM BODIPY-ATP, discrete single, fluorescent-nucleotide binding events were observed within myofibrils (Fig. 3A). We interpret these events as specific binding of a fluorescent ATP molecule, followed by hydrolysis and product release (fluorescent ADP). As in other studies, sarcomeric ATP binding is attributed to myosin, as it is the predominant ATPase in the sarcomere (Stewart et al., 2010). Individual event lifetimes were determined by kymograph analysis (Fig. 2A) and are a direct measure of the ATPase cycle time, given that the product release steps of the cycle are rate-limiting. Similar to our previous studies using the identical approach in slow skeletal muscle (Nelson et al., 2020), fluorescent ATP binding lifetime distributions were best fit as the sum of two exponential processes (Fig. 2B), with time constants of 25±4s and 161±11s (N=972 events from 6 mice). These time constants are similar to those reported in the literature for various striated muscles and thus indicative of the DRX and SRX states, respectively (Hooijman et al., 2011). However, cardiac myofibrils demonstrated a greater overall fraction (57±5%) of SRX events, similar to other reports for mouse cardiac muscle (Toepfer et al., 2020), but approximately twofold greater than skeletal muscle (Nelson et al., 2020; Stewart et al., 2009).

**Figure 3.**
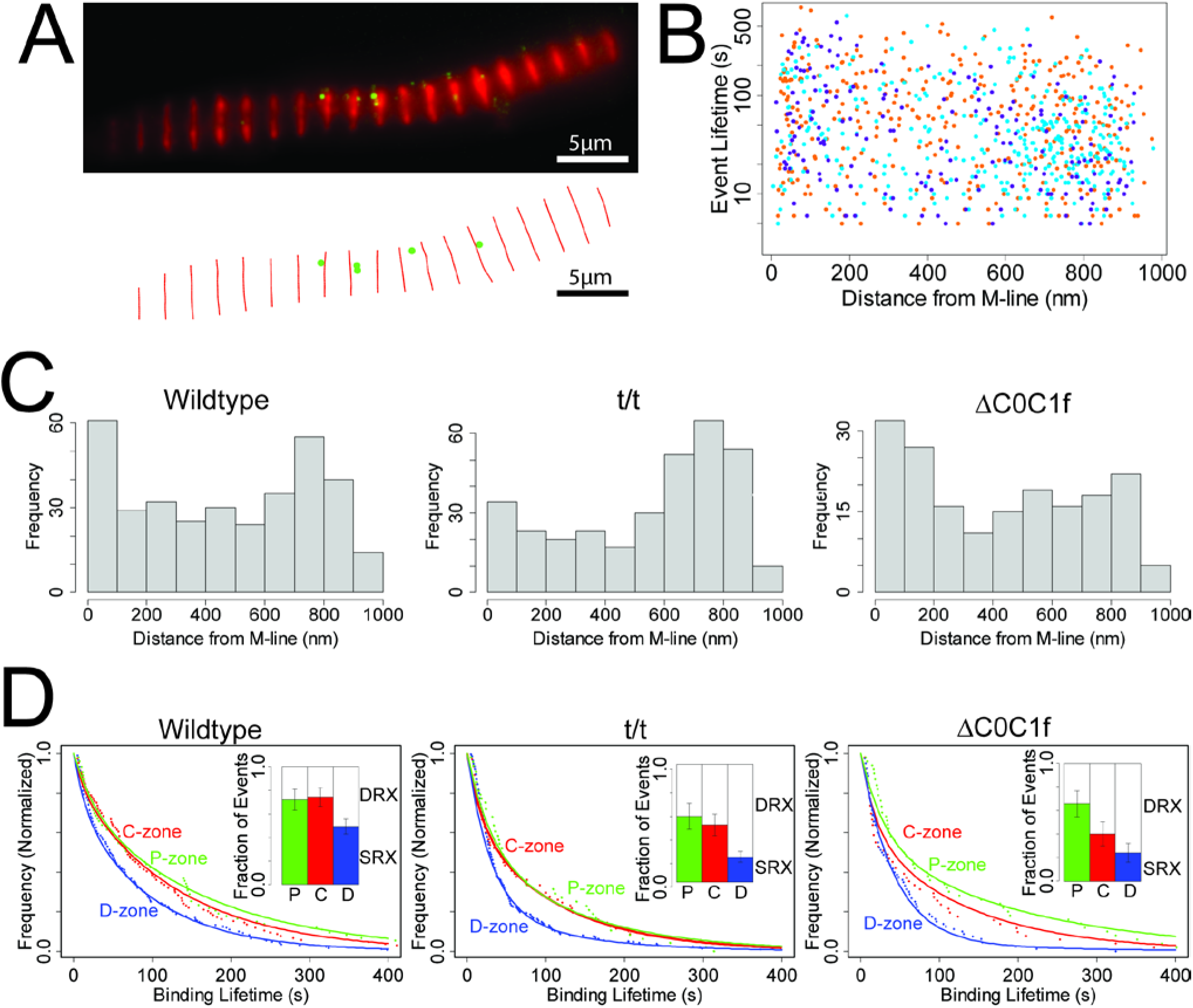
Event Localizations and Distributions. A) (upper) combined fluorescence image showing anti-myomesin (red) and BODIPY-ATP (green), combined from 100s of acquisition. (lower) X-Y plot of resulting localization data for same interval. B) Scatterplot showing distance from M-line and lifetime for each localized event. WT (orange), ΔC0C1f (purple), and t/t (blue). C) Histograms showing event frequency as a function of distance from the M-line. D) Survival plots showing difference for events localizing to C-zone (155-500nm from M-line) and D-zone (> 500nm from M-line). Insets are proportions of SRX and DRX events resolved by zones.

### Nucleotide Binding Event Locations

Utilizing super-resolution imaging methodologies, we determined the sub-sarcomeric location of each event (Fig. 3B). The well-defined sarcomeric geometry allowed us to assign each event to a sarcomere zone, based upon the distance to the nearest M-line, i.e., the sarcomere center. In mammalian striated muscle, the myosin-containing A-band is defined by the thick filament length; extending 800nm on either side of the M-line. Sarcomere lengths, as measured from M-line to M-line averaged 1.75±0.07μm (N=177 sarcomeres), and a combined spatial resolution of 21nm allowed us to parse each half of the A-band into 4 zones (Lee et al., 2015), specifically, the C-zone (155-500nm from M-line), comprised of both myosin and cMyBP-C, while the flanking P-zone (80-155nm from M-line) and D-zones (>500nm from M-line) are both devoid of cMyBP-C (Fig. 1B). Finally, the region nearest (<80nm) the M-line is referred to as the “bare zone” as this region does not contain myosin heads or MyBP-C. A plot of each event’s lifetime versus its distance from the M-line (Fig. 3B) shows no clear trend in lifetimes, but suggests elevated event frequencies near the M-line and thick filament tip (Fig. 3C). When binding events within WT myofibrils were binned according to their corresponding zones and their distributions fit with the double exponential model (Fig. 3D, Wildtype), we found that the P- and C-zones demonstrated nearly identical proportions of SRX events (74±8 (N=20) and 71±6% (N=177), respectively, p=0.43), significantly higher than the D-zone with 45±10% SRX (N=168; p=0.02), indicating a spatial differential in SRX location. Of the events that could be unambiguously assigned a sub-sarcomeric location, 85% localize to the myosin containing P-, C- or D-zones. Curiously, 13% of spatially resolved events (53/418 events) occurred in the bare zone. Given that this region is devoid of myosin heads, these events may arise from some other M-line localized ATPase, of which there are many (Lange et al., 2020).

### Deletion of cMyBP-C

Given that SRX myosin in the WT are enriched in the C-zone where cMyBP-C exists (Fig. 3D), we took advantage of an established cMyBP-C transgenic mouse line that is devoid of cMyBP-C (cMyBP-C^t/t^; McConnell et al., 1999) to determine if eliminating cMyBP-C alters the overall proportion of SRX and whether any positional bias for the remaining SRX still exists within the sarcomere. The cMyBP-C^t/t^ hearts were hypertrophied, consistent with prior reports (McConnell et al., 1999). Although myocytes from cMyBP-C^t/t^ mice have been reported to exhibit highly variable sarcomere lengths (Toepfer et al., 2019), the myofibrils used in our analysis demonstrated sarcomere lengths of 1.67±0.07μm (N=199 sarcomeres), not significantly different than WT (p=0.20). cMyBP-C^t/t^ myofibrils demonstrated a nearly 40% reduction in the overall SRX content compared to WT (35±5% vs. 57±5% SRX, respectively, p<0.001; N=882 events from 5 mice; Fig. 2B). This reduction in SRX was distributed across both the C- and D-zones (N=117 and 216, respectively), such that the C-zone continued to exhibit ~2-fold higher proportion of SRX myosin than the D-zone (49±9% vs. 22±5%, respectively; p = 0.02, Fig. 3D). The P-zone event lifetimes in cMyBP-C^t/t^ sarcomeres (N=24) were not different from those of the WT P-zone (p=0.57).

### N-Terminal cMyBP-C Truncation

The SRX reduction observed in the cMyBP-C^t/t^ myofibrils suggests that the presence of cMyBP-C contributes to SRX stabilization. If so, then are cMyBP-C N-terminal domains critical to this stabilization? We again took advantage of an existing cMyBP-C transgenic mouse line that expresses an N-terminal truncated cMyBP-C, lacking the C0 and C1 domains and the first 17 amino acids of the M-domain (cMyBP-C^ΔC0-C1f^; Lynch et al., 2021; Fig. 1C). As with the cMyBP-C^t/t^ null myofibrils, the cMyBP-C^ΔC0-C1f^ sarcomere lengths were no different than WT (1.75±0.06μm vs. 1.75±0.07μm, respectively, p=0.38). Also similar to the cMyBP-C^t/t^ myofibrils, the cMyBP-C^ΔC0-C1f^ myofibrils demonstrated a significant 45% reduction in the overall SRX content compared to WT (31±7% vs. 57±5%, respectively, p=0.02; N=469 events from 3 mice; Fig. 2B). This reduction in SRX occurred in both the C- and D-zones (N=101 and 80, respectively), such that the C-zone continued to exhibit ~2-fold higher proportion of SRX myosin compared to the D-zone (40±10% vs. 24±8%, respectively, p=0.20). Once again, the lifetime distribution of the P-zone events (Fig. 3D) was not significantly different than in the WT (N=25; p=0.93).

## Discussion

The SRX state appears to be common to all mammalian striated muscle myosin and may serve a significant role in modulating muscle contractility (Brunello et al., 2020). However, different muscle types show substantial differences in the overall proportion of SRX, evidenced by the twofold difference between the present data in mouse cardiac myofibrils compared to our previously published data (Nelson et al., 2020) in slow-twitch, rat soleus (57% vs. 28% SRX, respectively). The higher SRX content in cardiac tissue may represent a unique aspect of cardiac myosin. Unlike skeletal muscle, cardiac SRX myosin remain sequestered upon calcium activation (Hooijman et al., 2011), potentially acting as a “reserve pool” of myosin that can be recruited for additional contractile capability in response to physiological demand.

In both the slow-twitch, rat soleus (Nelson et al., 2020) and mouse cardiac myofibrils, SRX myosin are localized predominantly to the C-zone where MyBP-C resides. Where they differ is in the abundance of SRX in their D-zones, which are devoid of MyBP-C. Specifically, in the cardiac myofibril D-zone a significant SRX proportion (45%) are present compared to their minimal presence (9%) in the soleus. With the potential that myosin may exist in a dynamic equilibrium between SRX and DRX states (Nag and Trivedi, 2021), these data suggest this equilibrium may be shifted more towards the SRX state in mouse cardiac myosin than in skeletal muscle, perhaps enhanced within the context of the cardiac thick filament (Gollapudi et al., 2021). Additionally, this dynamic equilibrium appears to be further shifted by the existence of cMyBP-C in the C-zone, which in some manner stabilizes the SRX state such that it predominates within this zone.

If cMyBP-C stabilizes the SRX state, then eliminating cMyBP-C by genetic knock down should dramatically reduce the proportion of SRX, specifically in the C-zone. In the cMyBP-C^t/t^ null mouse, we did in fact observe a marked 40% reduction in the overall proportion of SRX myosin, including the expected reduction within the C-zone (Fig. 3D). The fact that a significant proportion of SRX remained in the C-zone (49%) of the cMyBP-C null supports the concept of a dynamic equilibrium between the SRX and DRX states being shifted inherently towards the SRX in cardiac muscle. This is also echoed by the similarity between the WT D-zone and the C-zone in the cMyBP-C^t/t^ null, as neither contains MyBP-C and both demonstrate ~45% SRX. However, the proportion of SRX in the D-zone of the cMyBP-C^t/t^ null was surprisingly lower than the WT D-zone. This reduction of SRX in the region of the sarcomere that is always devoid of cMyBP-C suggests that cMyBP-C may also act through some long range, indirect mechanism to stabilize the SRX state in the D-zone.

With cMyBP-C apparently stabilizing the SRX state, we asked whether this activity can be attributed to the N-terminal region, given its ability to modulate actomyosin contractile function both in *in vitro* motility (Razumova et al., 2006; Weith et al., 2012) and transgenic mouse studies (Lynch et al., 2021). Interestingly, myofibrils from the N-terminal truncated cMyBP-C^ΔC0-C1f^ mouse are almost indistinguishable from the cMyBP-C^t/t^ null, both in terms of the overall SRX proportion and SRX localization between the C- and D-zones (Fig. 3D). The similarity between the cMyBP-C^ΔC0-C1f^ and cMyBP-C^t/t^ null data emphasize the importance of cMyBP-C’s N terminus in stabilizing the SRX state. Interestingly, in a previous study (Lynch et al., 2021), we reported no difference in SRX content in ventricular muscle from the cMyBP-C^ΔC0-C1f^ and WT mice, as measured by mantATP chase experiments in muscle bundles. The apparent discrepancy with the present results may reflect a methodological difference, whereby tension applied to the muscle bundles (Lynch et al., 2021) may trigger a shift in the equilibrium towards the myosin DRX state (discussed below).

Perhaps most surprising is that a two-fold higher proportion of SRX exists in the C-zone of the cMyBP-C^t/t^ null compared to the D-zone. These data suggest that there must be an additional or alternative spatial or structural determinant that enriches SRX within the C-zone. For example, titin, which has super-repeats that differ in the C-zone and underlie the patterning of cMyBP-C into 11 stripes (Tonino et al., 2019) may contribute to stabilizing the SRX state, independent of cMyBP-C. Additionally, the structure of the thick filament backbone changes along its length, particularly within the D-zone as the thick filament tapers towards its tip (Sjostrom and Squire, 1977). In doing so, the landscape of myosin tails necessary for the folded SRX/IHM conformation, may present more opportunities for heads to be stabilized into the SRX state within the C-zone compared to the more distal thick filament D-zone.

What might be the functional consequence of such a SRX spatial gradient along the thick filament? Recently, a model of thick filament-dependent activation has been proposed (Linari et al., 2015) in which the SRX myosin contribute to the “off” state of the thick filament. Turning the thick filament “on” would then require that mechanical strain be sensed along the thick filament in order to trigger the release of myosin from the SRX state (Linari et al., 2015). Our data suggest that cardiac myosin motors are biased towards the DRX or “on” state within the D-zone, located at the ends of the myosin thick filament. These myosin would be the most available to bind the thin filament and generate force upon activation. Once bound, these myosin heads would cause strain to be distributed along the entire thick filament length (Ma et al., 2018), facilitating the release of more centrally located SRX myosin motors, particularly those that are enriched and stabilized by cMyBP-C in the C-zone. Therefore, with cMyBP-C having the capacity to stabilize the SRX state, future therapeutic design may target the cMyBP-C as a means of altering cardiac contractility.

## Acknowledgements

The authors would like to thank Mike Previs and Colleen Kelly (University of Vermont) for contribution of biological samples and Guy Kennedy (Instrumentation and Model Facility, University of Vermont) for technical expertise and imaging assistance, and James McNamara (Murdoch Children’s Research Institute) for insightful conversations.

